# Anatomical description of the jaw muscles and theoretical bite force assessment in South-American opossums using manual and virtual dissection methods

**DOI:** 10.1101/2025.06.12.659107

**Authors:** Melekian Alice, Decuypère Vincent, Herrel Anthony, Clarac François, Ladevèze Sandrine

**Author notes:** Corresponding and co-last author. The authors have participated equally in the design and the supervision of this study.

## Abstract

Marsupials (Marsupialia, Mammalia) represent a clade with highly varied ecologies. This is particularly true for opossums (South American marsupials), which are difficult to observe and collect. Consequently, few studies have focused on their bite force and the muscles of their masticatory apparatus, and there exists only scant information about the diet of certain species. Here we describe the masticatory muscles of several previously unstudied opossum species including *Caenolestes fuliginosus*, *Dromiciops gliroides,* and *Monodelphis touan.* We calculate the bite force of these species using data from both manual and virtual dissections and compare their theoretical bite forces with literature data. Additionally, we explore the differences between manual and virtual dissection to determine muscle PCSA (Physiological Cross-Sectional Area). We tried two virtual methods (VPDE: “virtual physiological data estimating method” and SM: “slicing muscle method”) to calculate the PCSA, determine the differences induced by the inter-fiber void in the virtual volume, and calculate a correction post-treatment with the contrast agent. The results highlighted variation in the position of the muscular attachments of the *M. zygomaticomandibularis*, whose insertion area is the largest in *Monodelphis touan* and the smallest in *Caenolestes fuliginosus*. The bite forces are coherent with estimates from the literature suggesting that the biomechanical model is reliable. The comparison between manual and virtual dissection showed that while virtual dissection allows an overall description of the masticatory muscles, it is more complex to accurately describe the different subdivisions of the muscle bundles. Virtual dissection data could potentially complete manual dissection data with the association of the VPDE method, the exclusion of inter-fiber voids, and a correction for the treatment with contrast-agents.

## Introduction

Marsupials (Marsupialia, Mammalia) are one of the three major clades of extant mammals, alongside monotremes (Monotremata) and placentals (Placentalia). Defined as the least inclusive clade containing *Didelphis marsupialis* Linnaeus, 1758, *Caenolestes fuliginosus* (Tomes, 1863), and *Phalanger orientalis* (Pallas, 1766) (Beck et al., 2014), Marsupialia comprises over 400 species (Eldridge, 2019; Mammal Diversity Database, 2024) that are distributed across seven clades occurring in the Americas and Oceania. Marsupials exhibit a remarkable ecological diversity including arboreal (*e.g.*, *Marmosa murina*), fossorial (*e.g.*, *Notoryctes typhlops*), semi-aquatic (*e.g.*, *Chironectes minimus*), gliding (*e.g.*, *Petaurus breviceps*), and terrestrial (*e.g.*, *Dasyurus viverrinus*) lifestyles. Their diets are equally diverse, encompassing carnivory (*e.g.*, *Sarcophilus harrisii*), insectivory (*e.g.*, *Marmosa murina*), herbivory (*e.g.*, *Macropus rufus*), and opportunistic omnivory (*e.g.*, *Didelphis virginiana*). However, the ecological and functional traits of many marsupials, particularly South American species, remain understudied.

Biting is a key functional trait allowing dietary diversification in different mammalian clades (*e.g.*, Aguirre et al., 2003; Santana et al., 2012; Cornette et al., 2015; Ginot et al., 2018; Kraus et al., 2022). In mammals, mastication involves several muscles—including the temporalis, the pterygoid, the masseter, the zygomaticomandibular, and the digastric—whose activation patterns vary during jaw opening and closing (Thexton & Hiiemae, 1975; Gorniak, 1985). These muscles form a morphological module, meaning that they are more functionally integrated with each other than with other body regions (Eble, 2005; Ziermann et al., 2021). The masticatory modules differ between marsupials and placentals and are associated with skeletal modules such as the neurocranium, which includes the squamosal, posterior cranial bones, mandible, and vertebrae (Ziermann et al., 2021). Morphological changes in masticatory muscles often influence skeletal structures and vice versa (Cornette et al., 2013; Fabre et al., 2018; Ferreira-Cardoso et al., 2020). Diet and feeding strategies are critical drivers of cranial and mandibular morphology as well as masticatory muscle architecture (Kienle et al., 2022; Bubadué et al., 2023). For example, dietary hardness influences masticatory muscle efficiency (Santana et al., 2010, 2012), while other adaptations, such as echolocation, also impact craniofacial structures (Jacobs et al., 2014; Giacomini et al., 2022 ; Takeuchi et al., 2024). Phylogeny plays an additional role, as distantly related species with similar ecological roles often exhibit divergent morphological adaptations to similar functional demands (Fabre et al., 2018; Ferreira-Cardoso et al., 2020). Moreover, marsupial development further constraints these adaptations, particularly due to the early onset of suckling behavior (Fabre et al., 2021). Additionally, social behaviors such as sexual selection and territoriality also influence bite force (Naretto et al., 2014; Thomas et al., 2015). *In vivo* studies have shown that bite force is to some degree plastic and variable over lifetime in response to both diet changes (Anderson et al., 2014) and behavioral choices regarding bite strength (Santana & Dumont, 2009). However, recent studies have shown that natural selection acts on variation in bite force (Herrel et al., 2016) and that bite force is heritable (Zablocki-Thomas et al., 2021).

Despite the global interest that has been given to the study of bite force in mammals, marsupials remain understudied in this context. This gap reflects a broader taxonomic bias in biological research, where marsupials are often sidelined or studied only for comparative purposes with placentals. Consequently, detailed descriptions of marsupial masticatory muscles remain fewer compared to those for placentals (but see Coues, 1872; Turnbull, 1967; Minkoff et al., 1979; Warburton 2009; Thomas et al., 2024; Decuypere et al. In press; Abreu & Astùa 2025). Variations in muscle nomenclature further complicate comparisons across studies, with marsupials diversity being frequently excluded from nomenclatural standardization efforts (Druzinsky et al., 2011; Diogo et al., 2016).

This study aims to start filling these gaps by examining the and masticatory muscle anatomy across various marsupial clades (Didelphimorphia, Paucituberculata and Microbiotheria). Here we provide new anatomical descriptions and bite force estimates for underrepresented species, particularly two South American marsupials: the colocolo opossum (*Dromiciops gliroides*), and the shrew opossum *Caenolestes fuliginosus*, for which dietary and muscular data remain scarce. This work integrates virtual dissections with contrast-enhanced CT-scanning to perform 3D-reconstruction of soft tissues and modeling of bite force to reduce reliance on physical specimens, extending its applicability to rare species (i.e., rare in natural history collections, and with understudied ecologies). As part of this effort, we further compare manual and virtual dissection techniques using two model opossum species, *Marmosa murina* and *Monodelphis touan*, to preliminary assess the consistency and reliability of virtual methods.

## Material and Methods

### Material

This study focuses primarily on small opossums, whose bite force and masticatory muscles remain poorly studied and largely undescribed. Our primary opossum model is *Marmosa murina*, two individuals of which having been manually dissected in a previous study (Decuypere et al., in press) and then scanned through X-ray microtomography.

*Marmosa murina (M1496,* adult female, collected in 2014) and *Monodelphis touan (M2838*, adult male, collected in 2017) were collected in the field during expeditions in French Guiana and are part of the JAGUARS (“Joindre l’Amazonie et la Guyane : Animaux, Ressources et Sciences », Institut Pasteur of French Guiana) collection. To complement this dataset and expand knowledge on the anatomy of rare South American taxa two specimens with available synchrotron data were included: *Caenolestes fuliginosus* and *Dromiciops gliroides* (Ladevèze *et al*., ESFR Experiment LS-2427 ED19/2015).

Finally, two skulls were added to the dataset: one from the Australian quoll (*Dasyurus viverrinus*), a marsupial cat that died at the MNHN zoo, and one from the *Philander opossum*. This improves phylogenetic coverage within Marsupialia. All specimens are listed in Table 1.

**Table 1:** Species, their inventory number, the sources of their dissection data, the scan resolution of their bone and the scan resolution of their soft tissues.

| Species | Inventory number | Dissection data | Scan resolution (bone) | Scan resolution (soft tissues) |
| --- | --- | --- | --- | --- |
| *Caenolestes fuliginosus* | MNHN.CG.1982.940 | Virtual | 6,23µm | 6,23µm |
| *Dasyurus viverrinus* |  | Thomas et al. (2024) | 40 µm |  |
| *Dromiciops gliroides* | MNHN.CG.1981.1075 | Virtual | 6,23 µm | 6,23µm |
| *Marmosa murina* | M1496 | Manual and virtual | 14µm | 14µm |
| *Monodelphis touan* | M2838 | Manual and virtual | 25µm | 13,4µm |
| *Philander opossum* | CG ZM.MO 2003.153 | Abreu and Astúa (2025) | 15µm |  |

The specimens were initially scanned using micro-computed tomography (micro-CT) at the AST-RX platform (UAR 2AD) of the MNHN, utilizing a "v|tome|x L240-180" ( Baker Hughes Digital Solutions). A first scan was performed to capture bone morphology. Afterward, the specimens underwent a chemical treatment (see section 2.2.1. Manual dissection method) to enhance the density and visibility of muscular tissues. They were then rescanned using CT imaging. Additionally, two ethanol preserved specimens from the collections of the MNHN were scanned at the Synchrotron (ESRF Grenoble) as part of a previous project.

### Nomenclature

All muscles involved in bite force production in marsupials were identified (*e.g.*, the masseter, zygomaticomandibularis, temporalis, and the medial and lateral pterygoid muscles). Muscle subdivisions had been previously identified in *Marmosa murina* (Decupere et al., in press). The nomenclature we used follows both Turnbull’s (1970) and the standardized *Nomina Anatomica Veterinaria* (5th edition; Waibl *et al*., 2012), adapted by Decuypere *et al*. (in press). The masseter complex is separated into pars superficialis, pars intermedius (depending on the species), and pars profundus. The temporalis muscle is subdivided into a pars zygomaticus, a pars superficialis, apars profundus lateralis and a pars profundus medialis. The zygomaticomandibularis muscle - often considered as part of the masseter complex - is a single bundle with an anterior and a posterior part. The medial and lateral pterygoid are made up of two single bundles. The digastric muscle, only involved in jaw opening, was excluded.

### Dissections

#### Manual dissection

Manual dissections were performed on one side of each specimen. All masticatory muscles, along with digastric muscles, tongue, and eye tissue, were removed from the dissected side. The skull and muscles from the non-dissected side were immersed in a 5% phosphomolybdic acid (PMA) solution diluted in 70% ethanol during three to nine weeks, depending on the impregnation time required for muscle staining. Dissected muscles were stored in 70% ethanol before being weighed. Weights were obtained using a METTLER AE 100 balance after blotting the muscles dry with a paper towel to remove the excess ethanol.

Individual muscles were submerged in a 30% aqueous nitric acid solution for 48 hours to dissolve connective tissues, following the method of Loeb and Gans (1986). After connective tissues were dissolved, the nitric acid was removed and replaced by a 50% aqueous solution of glycerol. Fibers were isolated using tweezers and photographed under a Leica WILD M3Z binocular microscope with oblique lighting (Intralux® 4000, Geprüfte Sicherheit) on a black background. The images were analyzed in ImageJ (Schneider *et al*., 2012).

#### Virtual dissection

After soaking in PMA, specimens were re-scanned. Virtual dissections were performed in Mimics 3D 23.1 (Materialise, Leuven, Belgium). The muscles were segmented slice by slice because grayscale thresholds were insufficient to isolate them. Manual outlining was done every 5–10 slices using Mimics’ interpolation tool. The segmentation was simultaneously processed in sagittal, coronal, and axial views, with a 3D rendering displayed in a separate window that allowed to monitor the modeling progress.

Muscles were segmented individually and separated from the remaining cranial structures using Boolean operations. Each muscle was segmented based on its observed morphology during manual dissection. 3D objects were created from segmented masks and exported as .stl files for further treatment in Geomagic Wrap (3DSystems, Morrisville, NC). In order to manage the file size, the mesh density was reduced to about one million polygons per object while maintaining sufficient resolution for measurements.

Muscle volume and fiber lengths were estimated by a first virtual method (named “**v**irtual **p**hysiological **d**ata **e**stimating method” or “**VPDE**”) using Mimics’ volume and measurement tools. The gap between the fibers was alternatively included and excluded (Figure 1) to explore the differences in muscle volume. Comparisons between manual and virtual PCSA estimates were made for specimen M1496 and M2838 with percentage differences calculated. Another method (named “**s**licing **m**uscle method” or “**SM**”) for estimating the PCSA (physiological cross section area; defined below) was tested for *Marmosa murina* (M1496) using the Geomagic software and the "estimate area" tool from the "analysis" tab. This tool allows a 3D object to be sliced along a plane whose position in space can be adjusted. The PCSA (in cm²) is obtained by slicing the largest cross-sectional area of the muscle perpendicular to the fiber direction. This effectively corresponds to the anatomical cross-sectional area of the muscles (ACSA) which is by definition lower than the PCSA for pennate muscles.

**Figure 1.**
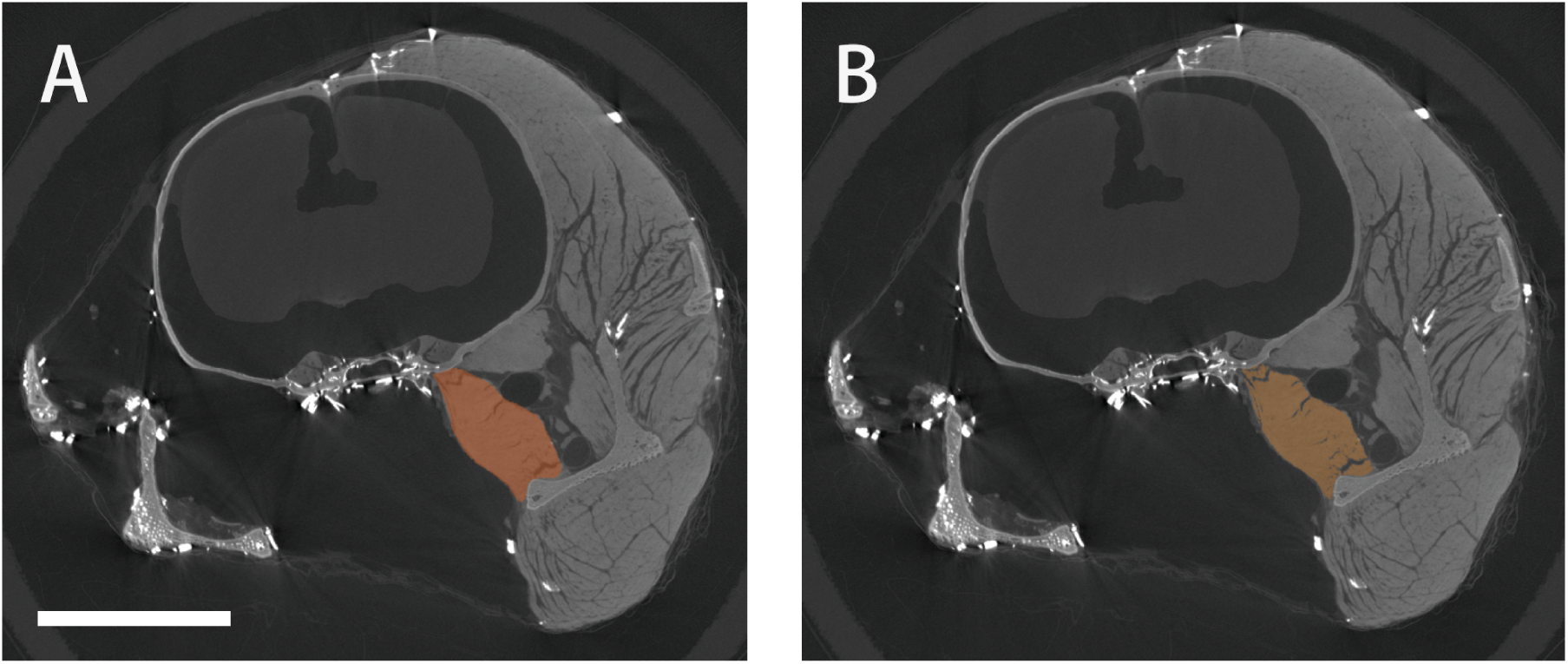
-. Differences on a slice between a segmentation with the gaps (A) and without (B) for the *M. pterygoideus medialis* of *Marmosa murina* (M1496). **Scale bar**: 5mm.

### Biomechanical model

#### Muscle force estimation

Bite force estimation first relies on calculating the theoretical maximal force that is generated by each masticatory muscle:

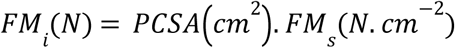

The PCSA is the physiological cross section area of the muscle. 𝐹𝑀_𝑠_ is the muscle specific tension which have a constant value of 30 *N.cm*^-1^ for mammals (Nigg et Herzog 1994)

PCSA was determined as:

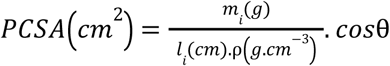

Where: *m_i_*: muscle mass; *l_i_*: fiber length of the muscle; ρ: muscle density which is a constant value of 1.056 *g. cm*^−3^ (Méndez and Keys, 1960; Leonard et al., 2020).

The pennation angle was excluded, consistent with previous studies (Ginot *et al*., 2018; Hartstone-Rose *et al*., 2018) because it has been argued to be more accurate to calculate PCSA without pennation angle when the multipennate muscles are too complex as found in jaw muscles (Hartstone-Rose *et al*., 2018; Martin *et al*., 2020; Hartstone-Rose *et al*., 2019). Correction factors were applied to estimate *in vivo* muscle mass based on preservation methods: 1.32 for formalin and 1.69 for 70% ethanol (Leonard *et al*., 2022a, 2022b). As the PMA increases the density of the muscles in order to enhance them, it also increases their volume. No corrections factors are available in literature to correct this effect. Considering that muscles and bones have similar threshold values (so similar density) after the treatment with the contrast agent, the correction factor named “**c**orrection **p**ost **c**ontrast **a**gent” was determined as:

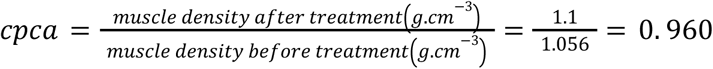

Where the muscle density after treatment is considered as the same as the mandibular bone density, a constant value of 1.1 *g . cm*^−3^(Devlin et al. 1998).

#### Anatomical Coordinate System (ACS) and bite force calculation

To project each muscle force onto the jaw-closing axis and calculate the theoretical bite force, an anatomical coordinate system was defined in Geomagic. The origin was set at the temporomandibular joint (TMJ) and the system is defined by three axes (*i.e*., the anteroposterior axis x, the mediolateral axis y, and the dorsoventral axis z), which enable the vectorial projection of each muscle force.

Each masticatory muscle is modeled as a vector connecting the centroid of the insertion area (IA) on the mandible to the centroid of the origin area (OA) on the skull. Each vector is projected onto the three axes, allowing the calculation of each muscle force using:

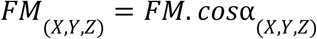

For the specimens whose muscles were segmented, the origin and insertion areas are defined by a Boolean intersection operation between the 3D model of the muscle and the 3D model of the skull or mandible. For the specimens that were manually dissected, the areas were manually traced based on muscle observations during dissections or available dissection data (Thomas et al., 2024; Abreu et Astùa 2025). The centroid of these areas is then calculated using the "centroid" tool in the "feature" tab. Finally, the various lever arms are determined with the "distance" tool in the "analysis" tab, by measuring the orthogonal projection of the distance between the centroid of each muscle IA and the ACS origin along the X axis (as schematized in Figure 2).

**Figure 2.**
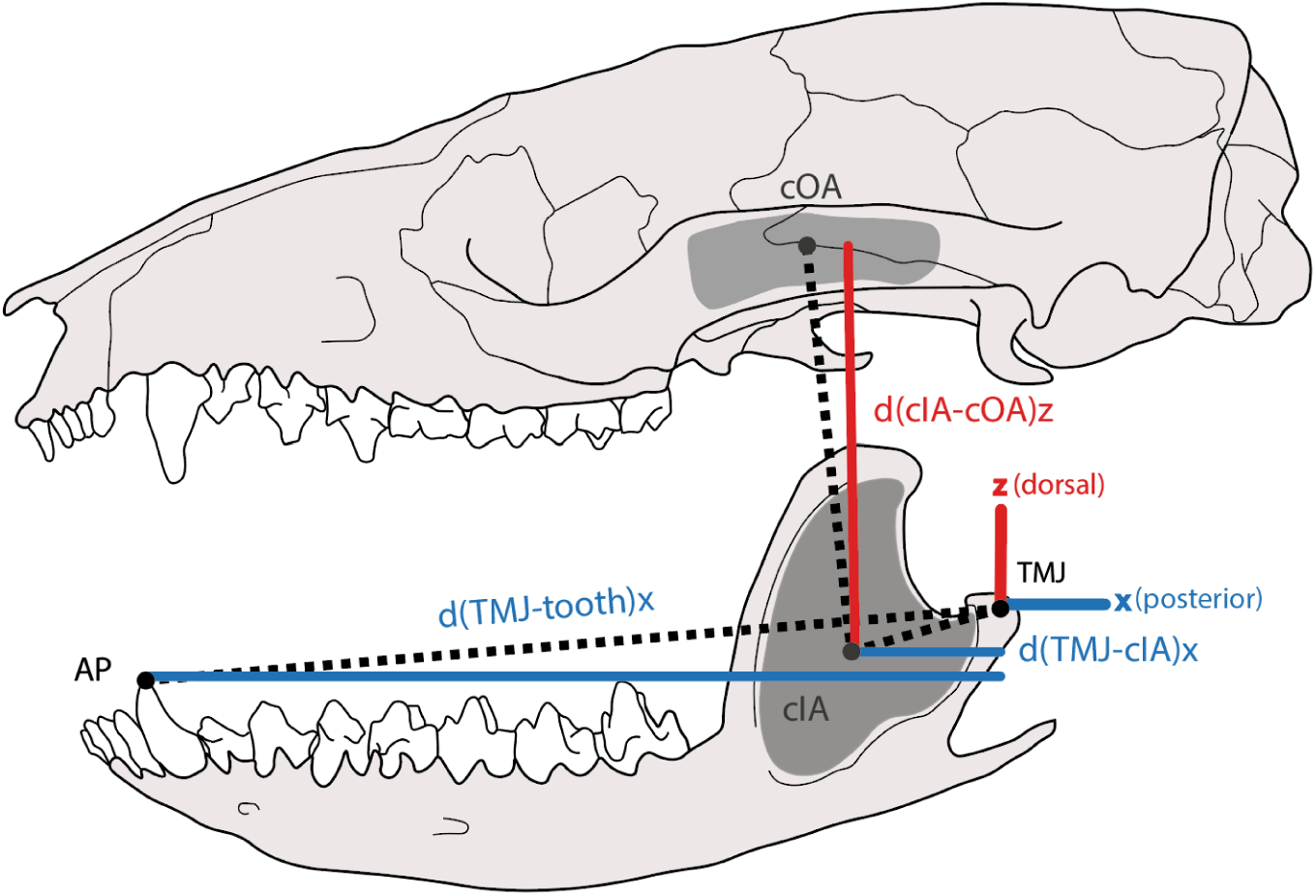
-. Figure showing the various distances used in the bite force calculation on the lateral view of the skull and mandible of *Monodelphis brevicaudata* (modified from Wible, 2003). **Legend: cIA**: Centroid of the insertion of the *M. zygomaticomandibularis*. **cOA**: Centroid of the origin of the *M. zygomaticomandibularis*. **TMJ**: Temporomandibular joint. **x**: Anteroposterior axis. **z**: Dorsoventral axis. **Distances:** Black, dashed line: Total distance between points. **Red**: Distance along the dorsoventral axis. **Blue**: Distance along the anteroposterior axis.

Each muscle in the system generates a moment Miz (N.cm⁻¹) that reflects its ability to rotate the mandible around the jaw’s articulation point (i.e., the origin of the ACS). This moment is the product of the vertical (z) component of the muscle force FM (in Newtons) and the lever arm d (in cm), which corresponds to the longitudinal distance (x) between the pivot point (TMJ) and the centroid of the muscle’s insertion area (IA). Moments were calculated as:

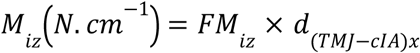

Where *FM_iz_* is the vertical component of the force, and *d_(TMJ-cIA)x_* is the lever arm length.

The theoretical bite force is calculated as the ratio between the sum of the moments ∑ *M_iz_* of the masticatory muscles and the out-lever *d_(TMJ-tooth)x_*, which corresponds to the longitudinal distance (x) between the pivot point (TMJ) and the tooth where the bite force is applied. To compute the bilateral bite force, this ratio is multiplied by two:

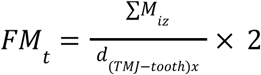

The theoretical bite forces were compared to results from other modeling studies: Abreu et Astùa (2025) and Wroe *et al*. (2005).

## Results

### Manual dissection (Raw data in Appendix 1)

*Description of the dissection of the masticatory muscles of* Monodelphis touan *(M2838) (Fig. 3):*

**Figure 3:**
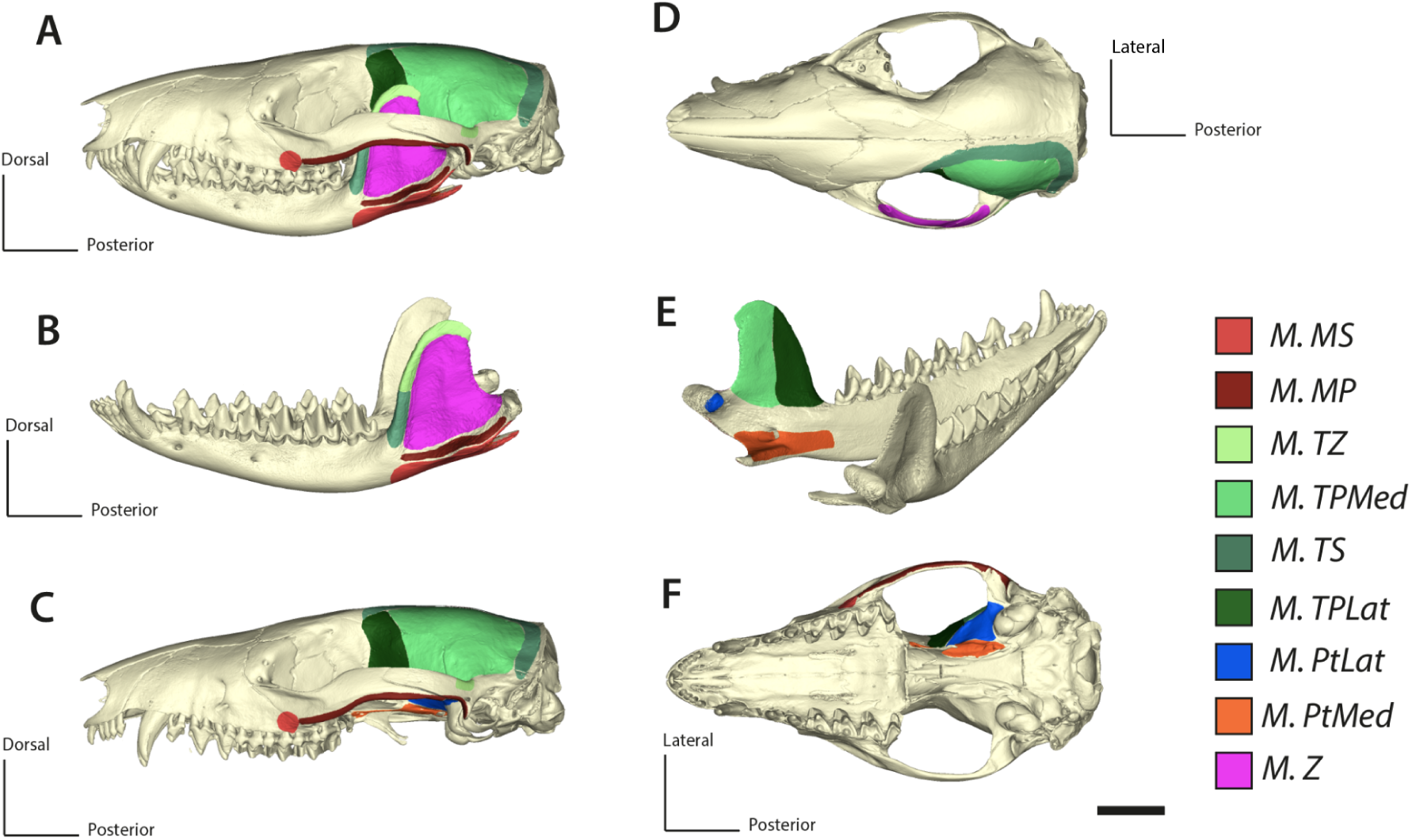
Results of the manual dissection of *Monodelphis touan* (M2838). The colored areas represent the different attachment areas (insertion area IA and origin area OA) of the masticatory muscles. Some of the muscles are attached by a tendon or a thin aponeurosis. **Abbreviations**: M.MP: *M. masseter profundus*. M.MS: *M. masseter superficialis*. M.PtLat: *M. pterygoideus lateralis*. M.PtMed: *M. pterygoideus medialis*. M.TPLat: *M. temporalis profundus lateralis*. M.TPMed: *M. temporalis medialis*. M.TS: *M. temporalis superficialis*. M.TZ: *M. temporalis zygomaticus*. M.Z: *M. zygomaticomandibularis*. **Scale**: 5mm

#### Digastric muscle (Di)

The digastric muscle accounts for approximately 4.98% of the total masticatory muscle mass. It consists of two parts, anterior and posterior, connected by a thick tendon. A thin aponeurosis covers the anterior part and thickens towards the posterior part. The muscle originates from the most posterior part of the exoccipital and inserts medioventrally on the mandibular body, from the lower third premolar (pm3) to the angular process. The anterior part is diamond-shaped, long, and flat, while the posterior part is thicker and tapered. The tendon connecting the two parts is located medioventrally to the superficial masseter muscle. The muscle fibers are oriented anteroposteriorly throughout.

#### M. masseter superficialis (MS)

The superficial masseter muscle accounts for approximately 17.97% of the total masticatory muscle mass. A thick aponeurosis clearly separates it from the digastric muscle and covers its entire surface. The muscle is thick and round, originating from a thick tendon located at the lateral posterior part of the maxilla above the posterior section of the second upper molar (M2). It inserts ventrolaterally on the mandibular ramus along the masseteric fossa, from the posterior third lower molar to the angular process, and extends upward to the condylar process. The muscle is pennate.

#### M. masseter profundus (MP)

The deep masseter muscle represents approximately 6.19% of the total masticatory muscle mass. It has a rounded and flat appearance, originating from the lower lateral ventral part of the zygomatic arch and inserting ventrally in the masseteric fossa, from its most posterior part to almost the third lower molar (M3). Its fibers are dorsoventrally oriented.

#### M. zygomaticomandibularis (ZM)

The zygomaticomandibularis muscle represents approximately 20% of the total masticatory muscle mass. It consists of two parts, anterior and posterior. It originates along the entire medial edge of the zygomatic arch and inserts across the masseteric fossa, above the insertion of the deep masseter muscle. Its fibers are loosely connected, with less connective tissue than in other masticatory muscles.

#### M. temporalis pars suprazygomatica (TZ)

The temporalis pars suprazygomatica accounts for 5.71% of the total masticatory muscle mass. It is external to the superficial temporalis muscle. It originates from the posterior part of the zygomatic arch, extending along half of its dorsal surface as a thin muscle layer. It inserts dorsolaterally on the upper half of the coronoid process.

#### M. temporalis superficialis (TS)

The superficial temporalis muscle constitutes 14.11% of the total masticatory muscle mass. It is covered by a thick aponeurosis that extends over half of the lateral surface of the zygomatic arch. It inserts dorsolaterally on the lower half of the coronoid process and originates along the sagittal and nuchal crests, bordering the frontal, parietal, interparietal, and squamosal bones.

#### M. temporalis profundus lateralis (TPLat)

The lateral deep temporalis muscle accounts for 4.04% of the total masticatory muscle mass. It is difficult to distinguish from the medial deep temporalis muscle due to the presence of abundant connective tissue. The muscle originates from the posterior part of the supraorbital margin of the frontal bone, extending from the sagittal crest to the frontal-palatal suture. It inserts anteriorly on the medial side of the coronoid process.

#### M. temporalis profundus medialis (TPMed)

The medial deep temporalis muscle is the most massive masticatory muscle, accounting for 29.72% of the total masticatory muscle mass. It originates posteriorly from the lateral region of the frontal bone, covering parts of the parietal, alisphenoid, and squamosal bones. It inserts medially on the coronoid process, posterior to the insertion of the lateral deep temporalis muscle.

#### M. pterygoideus lateralis (PtLat)

The lateral pterygoid muscle represents 0.97% of the total masticatory muscle mass, making it the least massive masticatory muscle. It originates ventrolaterally on the alisphenoid and inserts on the medial border of the mandibular condyle.

#### M. pterygoideus medialis (PtMed)

The medial pterygoid muscle constitutes 5.98% of the total masticatory muscle mass. It is long and flat, originating from the ventrolateral edge of the pterygoid and posterior part of the palatine. It inserts on the medial side of the mandibular ramus, bordering the insertion of the superficial masseter, from the angular process to the posterior part of the third lower molar. It also inserts into the pterygoid pit.

### Virtual dissection (Raw data in Appendix 2)

#### Marmosa murina, Caenolestes fuliginosus *and* Dromiciops gliroides

Virtual dissection of *Marmosa murina* (M1496) allowed the individualization of the masseter muscle into its superficial, intermediate, and deep parts, which was difficult to do in the manual dissection (Decuypere et al., in press; Fig. 4). However, it did not allow the accurate separation of the fascicles of the temporal muscle, except for the *M. temporalis zygomaticus*. The remaining temporal layers were grouped into a compact mass, making detailed fascicle separation challenging. The aponeurosis and tendons were hard to identify, making the attachment areas less extensive compared to the manual dissection.

**Figure 4:**
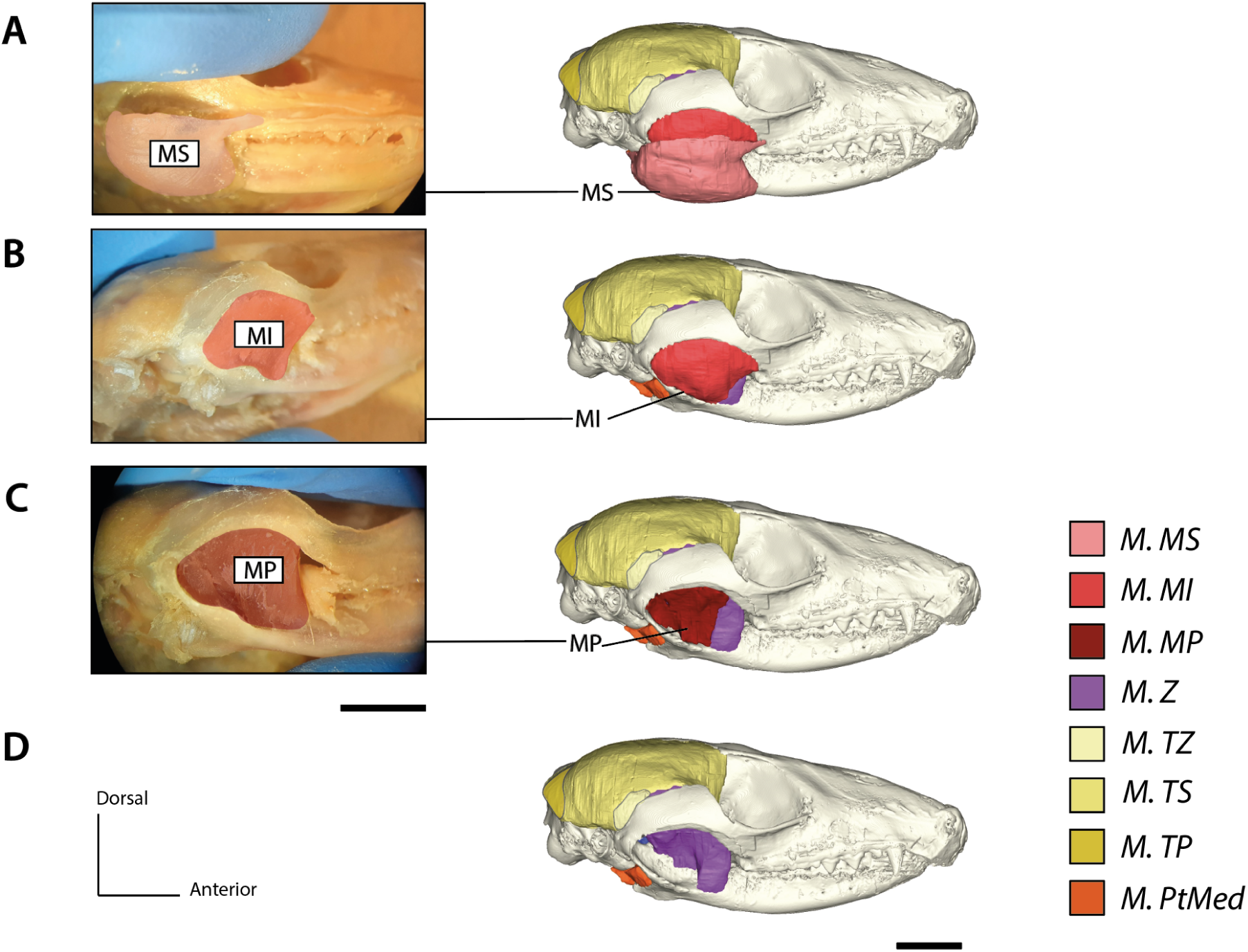
Results of the manual dissection and virtual dissection of the *M. masseter* different layers from *Marmosa murina* (M1496). **Abbreviations**: M.MI: *M. masseter intermedius*. M.MP: *M. masseter profundus*. M.MS: *M. masseter superficialis*. M.PtLat: *M. pterygoideus lateralis*. M.TP: *M. temporalis profundus*. M.TS: *M. temporalis superficialis*. M.TZ: *M. temporalis zygomaticus*. **Scale**: 5mm

Virtual dissections of *Caenolestes fuliginosus* (MNHN.CG.1982.940) and *Dromiciops gliroides* (MNHN.CG.1981.1075) provided initial insights into their masticatory muscles, previously undescribed due to the rarity of these species. These dissections were less precise than those of *Marmosa murina* but allowed the first estimates of the PCSA and attachment areas for their masticatory muscles using Boolean intersection operations.

*Masticatory muscle description of* Caenolestes fuliginosus (MNHN.CG.1982.940) (Fig. 5)

**Figure 5:**
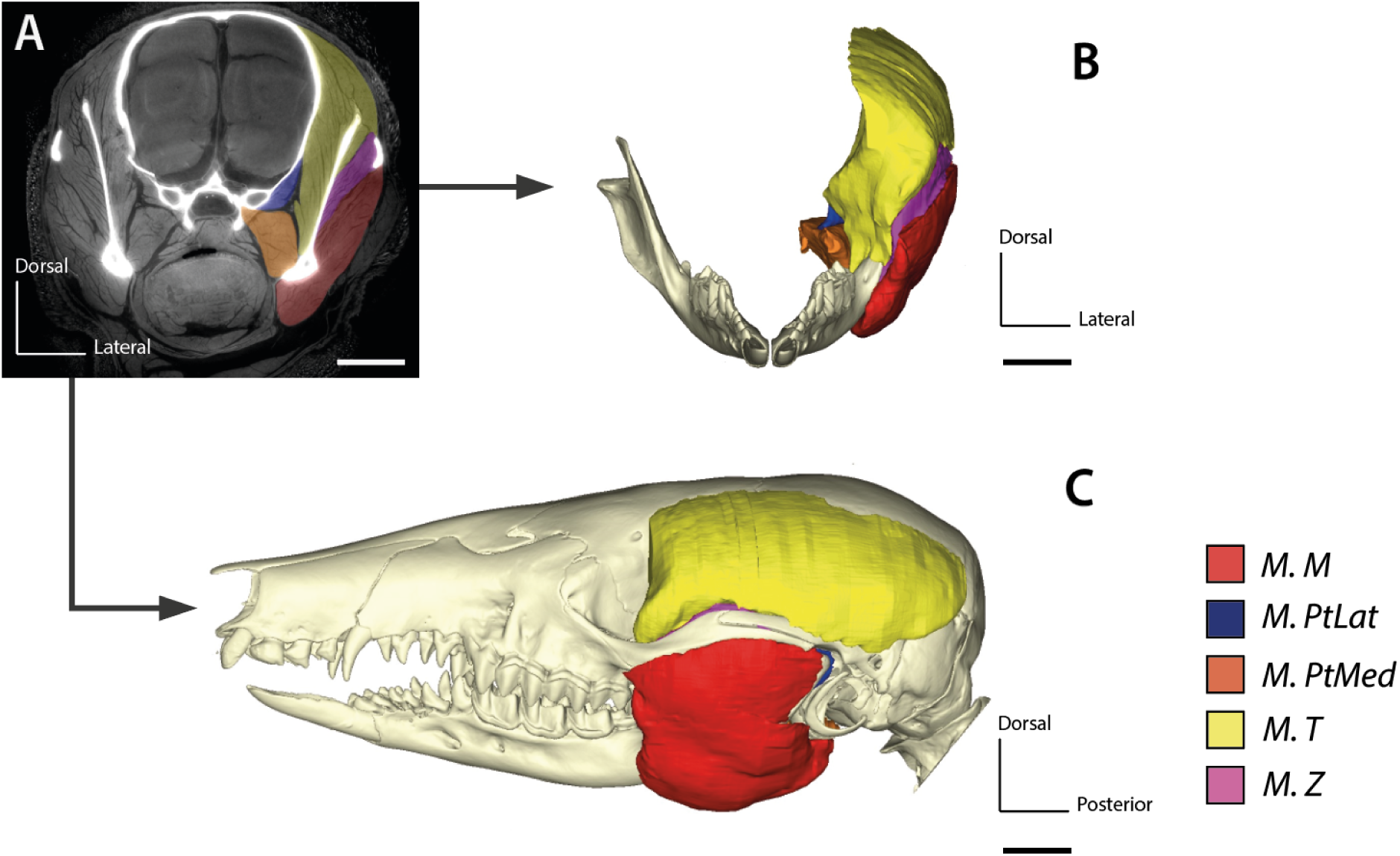
Results of the virtual dissection of *Caenolestes fuliginosus* (MNHN.CG.1982.940). **Muscles**: M.M: *M. Masseter*. M. PtLat: *M. pterygoideus lateralis*. M.PtMed: *M. pterygoideus medialis*. M.T: *M. temporalis*. M.Z: *M. zygomaticomandibularis.* **Views**: A: Coronal view of the CT image. B Coronal view of the 3D model. C Lateral view of the 3D model. **Scale bar**: 5mm

**Figure 6:**
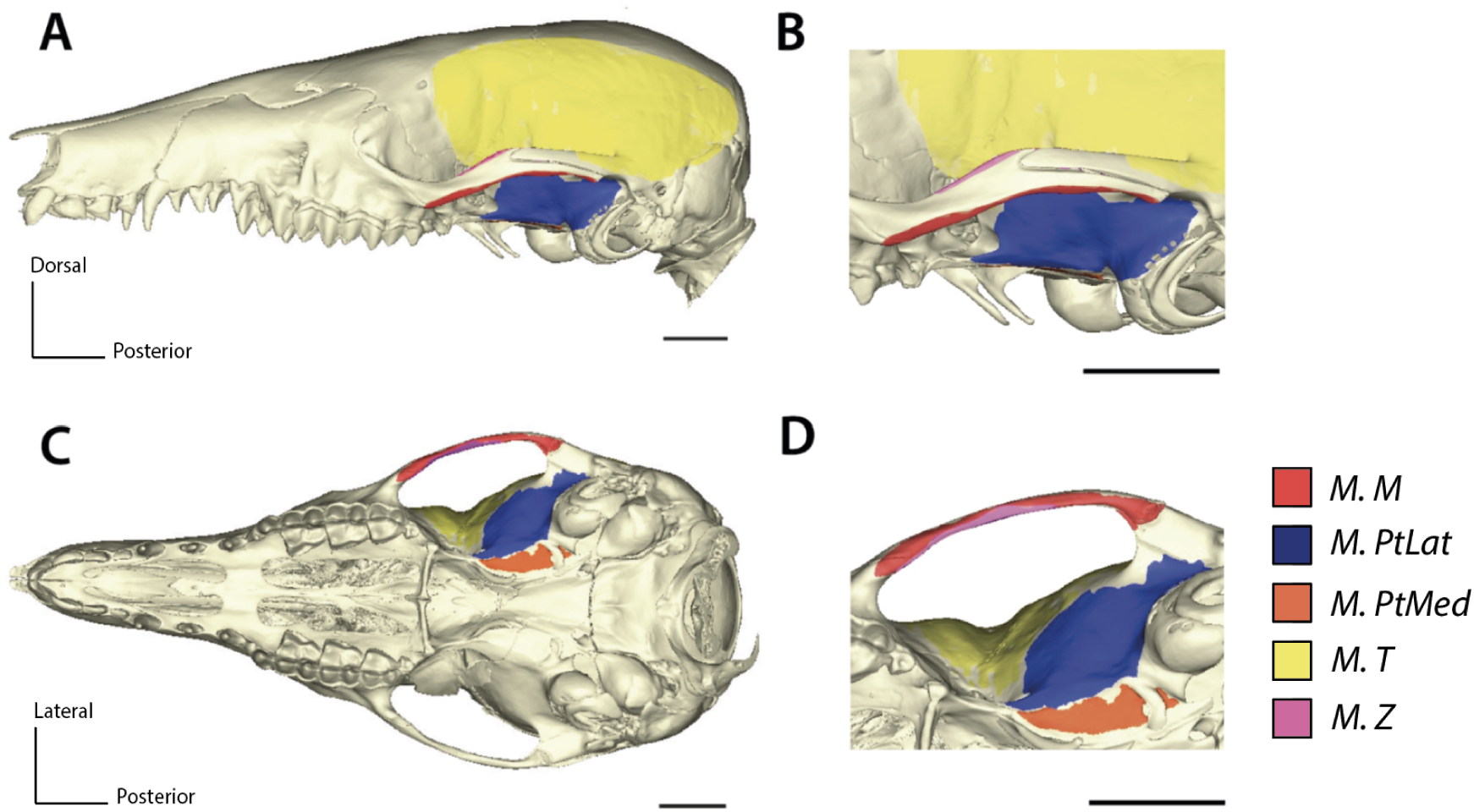
Results of the virtual dissection of *Caenolestes fuliginosus* (MNHN.CG.1982.940) and the origin areas (OA). **Origin areas**: M.M: *M. Masseter*. M. PtLat: *M. pterygoideus lateralis*. M.PtMed: *M. pterygoideus medialis*. M.T: *M. temporalis*. M.Z: *M. zygomaticomandibularis.* **Views**: A: Lateral view of the cranium. B: Lateral view, zoom on the zygomatic arch. C: Ventral view of the skull. D: Ventral view, zoom on the interorbital fossa. **Scale bar**: 5mm

#### M. masseter (M)

The masseter muscle accounts for 25.25% of the total muscle mass. It originates from the posterolateral portion of the maxilla, above the posterior part of the third upper molar, and extends along the ventral edge of the zygomatic arch. It inserts onto the mandibular ramus in the posterior half of the masseteric fossa and along its border, from the condylar process to the most anterior edge of the fossa, passing through the angular process.

#### M. zygomaticomandibularis (ZM)

The zygomaticomandibularis muscle constitutes 7.15% of the total muscle mass. It originates from almost the entire medial surface of the zygomatic arch, as well as the dorsolateral surface of the jugal bone via a thin muscular band. It inserts on the anterior half of the masseteric fossa and laterally on the lower half of the coronoid process.

#### M. temporalis (T)

The temporalis muscle represents 54.88% of the total muscle mass. It originates below the sagittal crest, and continues to the nuchal crest. It also originates from the posterior part of the zygomatic arch. The origin extends over parts of the parietal, frontal, squamosal, interparietal, alisphenoid, and palatine bones, reaching the palatine region of the interorbital fossa. It inserts onto the medial surface of the coronoid process, extending to the edge of the pterygoid pit.

#### M. pterygoideus medialis (PtMed)

The medial pterygoid muscle accounts for 9.01% of the total muscle mass. It originates along the entire ventral surface of the pterygoid bone. It inserts into the pterygoid pit and on the medial side of the angular process.

#### M. pterygoideus lateralis (PtLat)

The lateral pterygoid muscle accounts for 3.69% of the total muscle mass. It originates from the alisphenoid bone, extending to the boundary with the glenoid fossa, and inserts on the medial side of the condylar process.

Dromiciops gliroides (MNHN.CG.1981.1075; Fig. 8)

**Figure 7:**
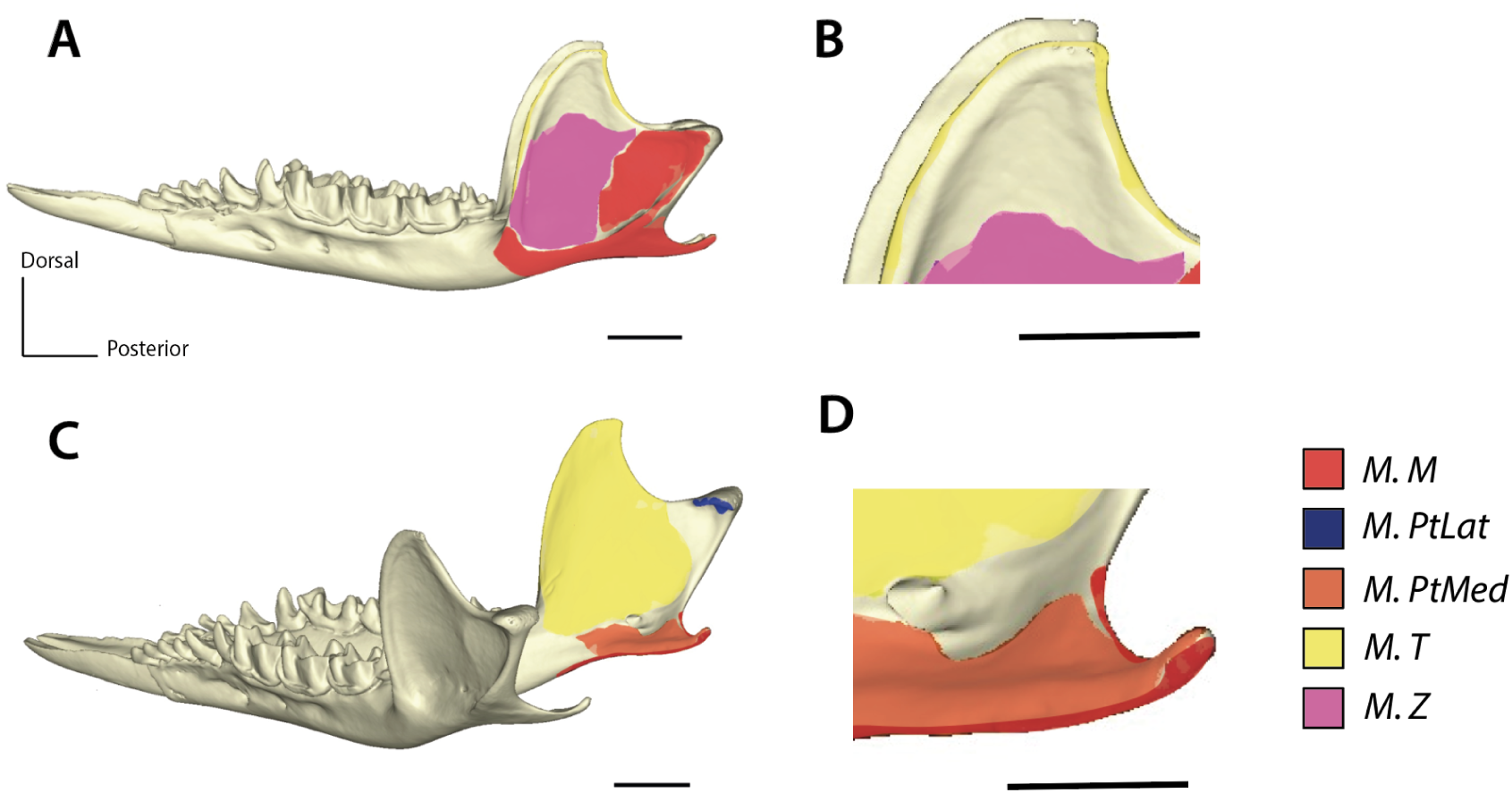
Results of the virtual dissection of *Caenolestes fuliginosus* (MNHN.CG.1982.940) and the insertion areas (IA). **Insertion areas**: M.M: *M. Masseter*. M. PtLat: *M. pterygoideus lateralis*. M.PtMed: *M. pterygoideus medialis*. M.T: *M. temporalis*. M.Z: *M. zygomaticomandibularis.* **Views**: A: Lateral view of the cranium. B: Lateral view, zoom on the zygomatic arch. C: Ventral view of the skull. D: Ventral view, zoom on the interorbital fossa. **Scale bar**: 5mm

**Figure 8:**
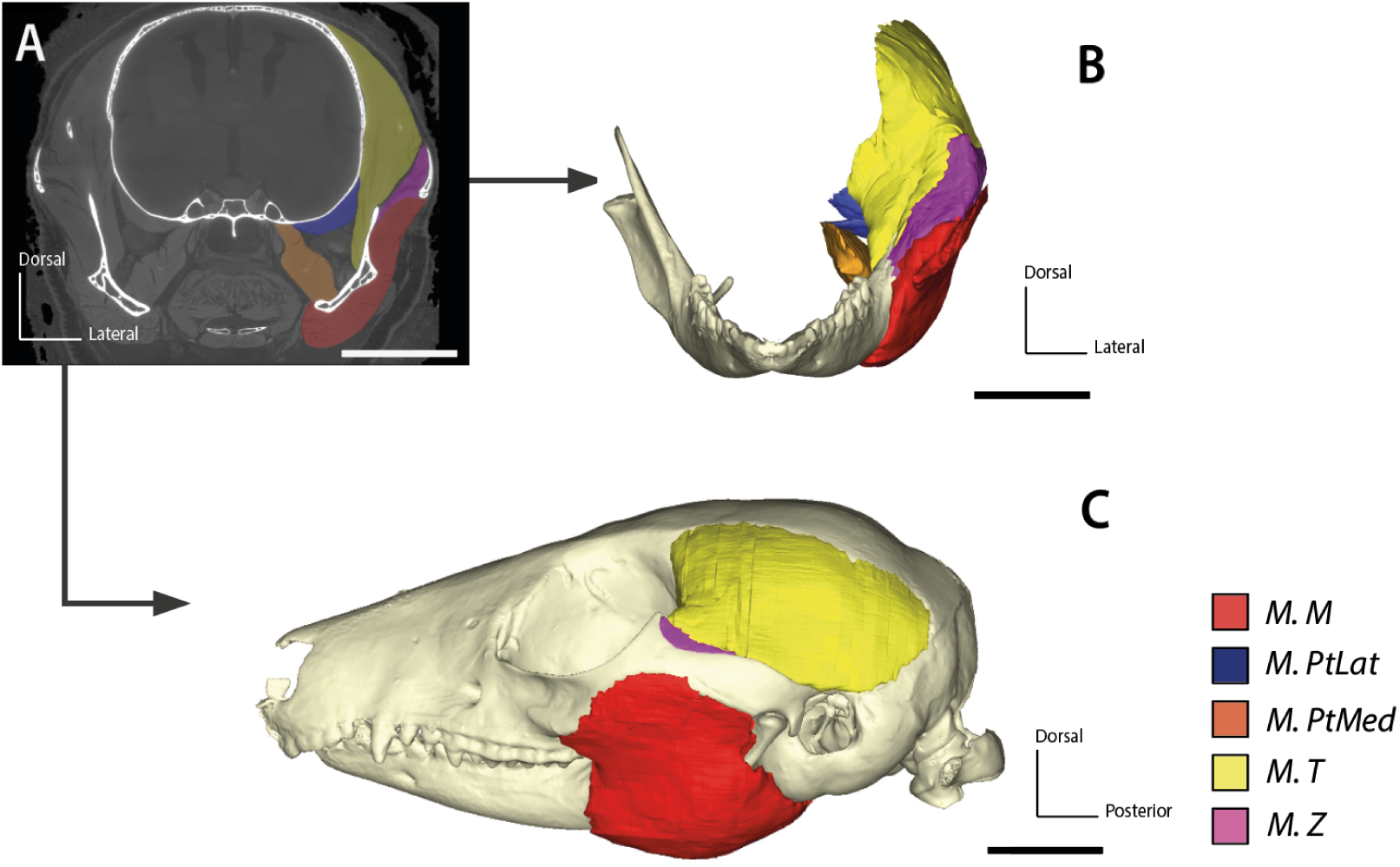
Results of the virtual dissection of *Dromiciops gliroides* (MNHN.CG.1981.1075). **Muscles**: M.M: *M. Masseter*. M. PtLat: *M. pterygoideus lateralis*. M.PtMed: *M. pterygoideus medialis*. M.T: *M. temporalis*. M.Z: *M. zygomaticomandibularis.* **Views**: A: Coronal view of the CT image. B: Coronal view of the 3D model. C: Lateral view of the 3D model. **Scale bar**: 5mm

**Figure 9:**
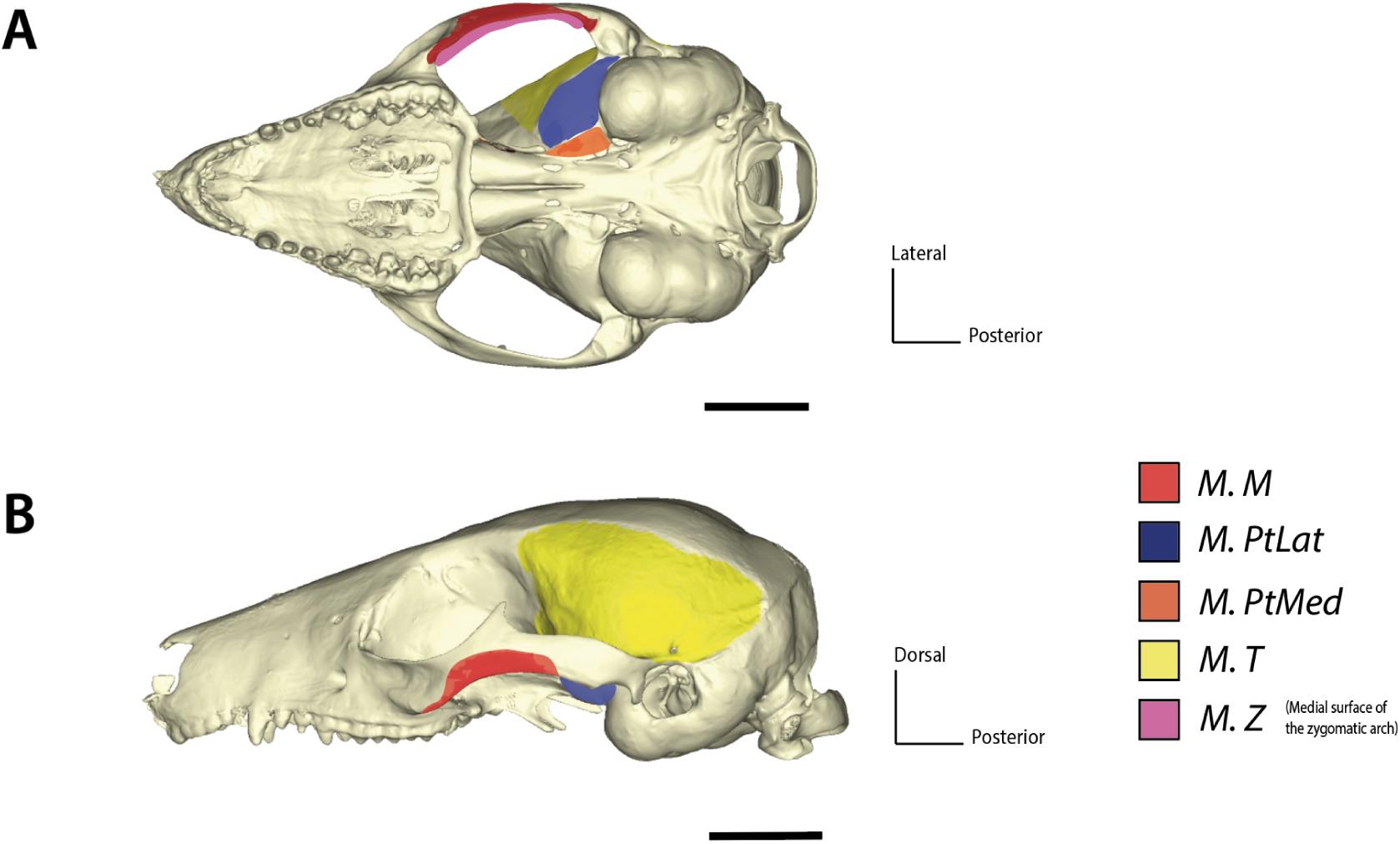
Results of the virtual dissection of *Dromiciops gliroides* (MNHN.CG.1981.1075) and the origin areas (OA). **Origin areas**: M.M: *M. Masseter*. M. PtLat: *M. pterygoideus lateralis*. M.PtMed: *M. pterygoideus medialis*. M.T: *M. temporalis*. M.Z: *M. zygomaticomandibularis.* **Views**: A: Ventral view of the cranium. B: Lateral view of the skull. **Scale bar**: 5mm

**Figure 10:**
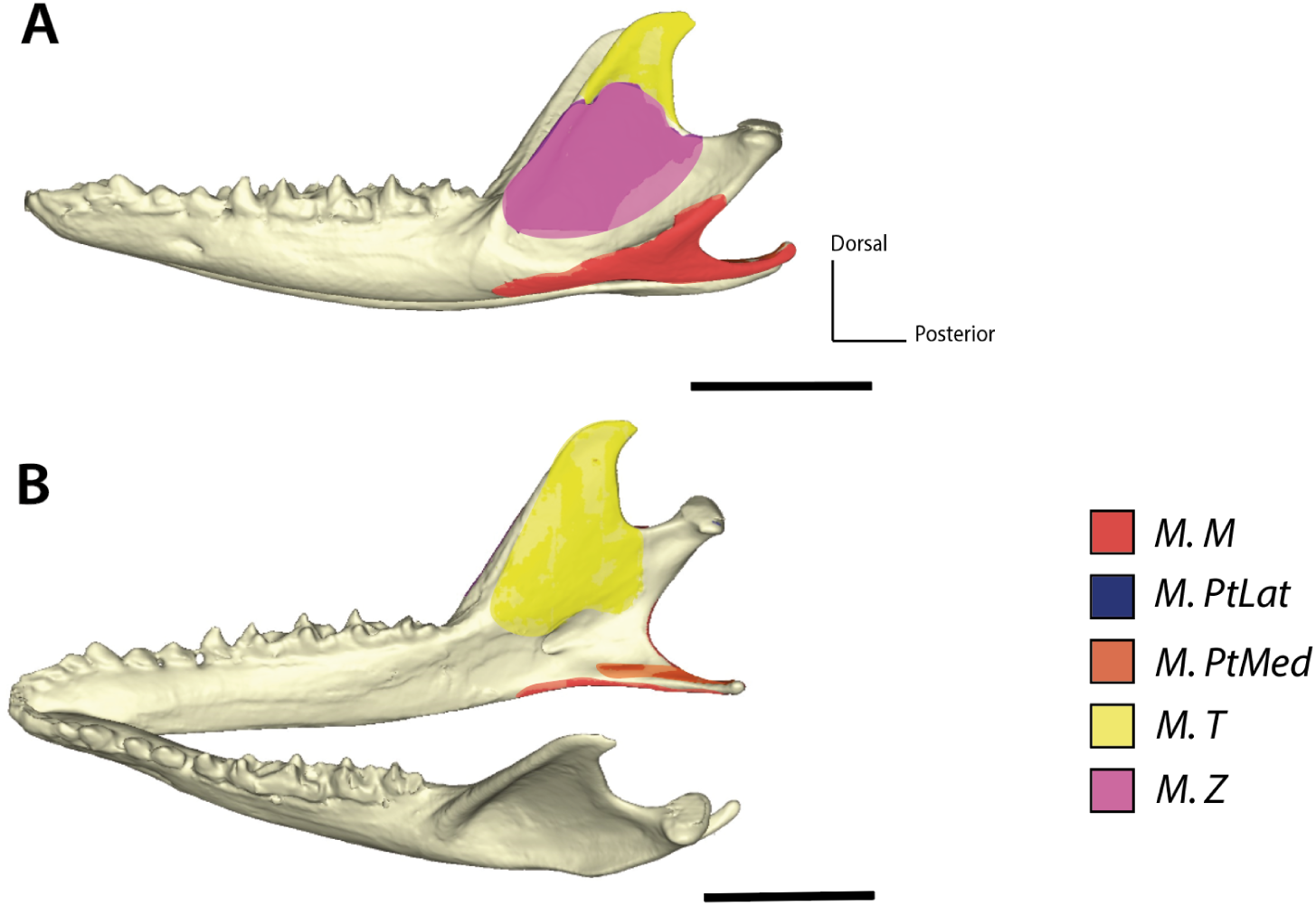
Results of the virtual dissection of *Dromiciops gliroides* (MNHN.CG.1981.1075) and the insertion areas (IA). **Insertion areas**: M.M: *M. Masseter*. M. PtLat: *M. pterygoideus lateralis*. M.PtMed: *M. pterygoideus medialis*. M.T: *M. temporalis*. M.Z: *M. zygomaticomandibularis.* **Views**: A: Lateral view of the mandible. B: Oblique view of the mandible. **Scale bar**: 5mm

*Dromiciops gliroides* differs from *Caenolestes fuliginosus* primarily in the distribution of muscle mass within the masticatory apparatus. Its *M. masseter* accounts for 32.98% of the total muscle mass, the *M. temporalis* for 39.54%, the *M. zygomaticomandibularis* for 11.9%, the *M. pterygoideus lateralis* for 3.39%, and the *M. pterygoideus medialis* for 12.19%.

The insertion of the *M. masseter* remains confined to the edge of the masseteric fossa and does not extend to the end of the condylar process. The origin of the *M. temporalis* on the zygomatic arch is much more posterior compared to that in *Caenolestes fuliginosus*. The insertion of the *M. temporalis* extends further down laterally on the coronoid process. The *M. zygomaticomandibularis* does not originate from the anterior dorsal portion of the zygomatic arch. It also does not originate ventrally on the zygomatic arch but remains confined to its medial surface.

### Comparison of PCSA and bite forces obtained using different methods

#### PCSA

The different methods of PCSA estimation (VPDE and SM) are summarized in Table 2 and Table 3. The margin of error between the VPDE method and the manual dissection estimation ranges from -0.33% to 33.01% for *Marmosa murina* and *Monodelphis touan*, depending on the muscle. The margin of error between the PCSA estimated for *Marmosa murina* with the SM method and that estimated by manual dissection ranges from -5.94% to 42.33%. Consequently, only the estimation performed with the VPDE method will be retained for specimens that could not be manually dissected, to minimize the error.

**Table 2:** Comparison of the different methods used to estimate the PCSA (cm²) of *Marmosa murina* including the percentage difference between the virtual and manual methods.

| M1469 | PCSA (cm<sup>2</sup>) |  |  |  |  |
| --- | --- | --- | --- | --- | --- |
| Muscles | PCSA manual dissection | PCSA virtual dissection<br>*No void*<br>*cpca correction*<br>*VPDE method* | %Difference with manual dissection | PCSA virtual dissection<br>*SM method* | %Difference with manual dissection |
| Masseter | 0.226 | 0.225 | (-)0.33% | 0.217 | (-)4.32% |
| Temporal | 0.355 | 0.345 | (-)2.80% | 0.335 | (-)5.94% |
| ZM | 0.077 | 0.072 | (-)7.41% | 0.129 | (+)40.24% |
| PtLat | 0.015 | 0.015 | (+)0.93% | 0.025 | (+)42.33% |
| PtMed | 0.125 | 0.130 | (+)3.84% | 0.095 | (-)31.85% |

**Table 3:** Comparison of the different methods used to estimate the PCSA (cm²) of *Monodelphis touan* including the percentage difference between the virtual and manual methods.

| M2838 | PCSA (cm<sup>2</sup>) |  |  |
| --- | --- | --- | --- |
| Muscles | PCSA manual dissection | PCSA virtual dissection<br>*No void*<br>*cpca correction*<br>*VPDE method* | % Difference |
| Masseter | 0.242 | 0.286 | (+)15.31% |
| Temporal | 0.487 | 0.539 | (+)9.63% |
| ZM | 0.050 | 0.057 | (+)12.04% |
| PtLat | 0.0185 | 0.028 | (+)33.01% |
| PtMed | 0.102 | 0.077 | (-)33.52% |

#### Masticatory muscles volumes

The volumes measured using virtual dissection are most of the time higher than those measured through manual dissection, except for the *M. pterygoideus medialis* of *Monodelphis touan*.

The difference between the results ranges under 20% (Table 5 and 6) for the *M. masseter*, *M. temporalis* and *M. zygomaticomandibularis* in the virtual dissection with all the corrections. It increases for the *M. pterygoideus medialis* and *M. pterygoideus lateralis (*up to 54.02%, Table 5 and 6).

**Table 4 -.** Comparison table of the different methods used to estimate the volume (µm^3^) of *Marmosa murina* muscles including the percentage difference between the virtual and manual methods and the different corrections.

| **M1469** | **Volume ($\text{mm}^3$)** |  |  |  |  |  |  |
| --- | --- | --- | --- | --- | --- | --- | --- |
| **Muscles** | **Volume in manual dissection** | **Volume in virtual dissection**<br>*No void*<br>*cpca correction*<br>*VPDE method* | **% Difference with manual dissection** | **Volume in virtual dissection**<br>*With void*<br>*cpca correction*<br>*VPDE method* | **% Difference with manual dissection** | **Volume in virtual dissection**<br>*With void*<br>*no correction*<br>*VPDE method* | **% Difference with manual dissection** |
| **Masseter** | 104.05 | 116.47 | (+)10.66% | 182.88 | (+)43.10% | 190.50 | (+)45.38% |
| **Temporal** | 194.32 | 228.22 | (+)14.85% | 328.55 | (+)40.85% | 342.24 | (+)43.22% |
| **ZM** | 39.93 | 45.42 | (+)12.08% | 85.94 | (+)53.54% | 89.52 | (+)55.40% |
| **PtLat** | 2.98 | 6.49 | (+)54.02% | 9.62 | (+)68.98% | 10.02 | (+)70.22% |
| **PtMed** | 26.43 | 31.00 | (+)14.70% | 51.82 | (+)48.97% | 53.97 | (+)51.01% |

**Table 5 -.** Comparison table of the different methods used to estimate the volume (µm^3^) of *Monodelphis touan* muscles including the percentage difference between the virtual and manual methods and the different corrections.

| **M2838** | **Volume ($\text{mm}^3$)** |  |  |  |  |  |  |
| --- | --- | --- | --- | --- | --- | --- | --- |
| **Muscles** | **Volume in manual dissection** | **Volume in virtual dissection**<br>*No void*<br>*cpca correction*<br>*VPDE method* | **% Difference with manual dissection** | **Volume in virtual dissection**<br>*With void*<br>*cpca correction*<br>*VPDE method* | **% Difference with manual dissection** | **Volume in virtual dissection**<br>*With void*<br>*no correction*<br>*VPDE method* | **% Difference with manual dissection** |
| **Masseter** | 147.4 | 167.44 | (+)12.0% | 214.397568 | (+)27.41% | 223.3308 | (+)34.01% |
| **Temporal** | 276.4 | 290.68 | (+)4.9% | 414.613901 | (+)29.60% | 431.88948 | (+)36.00% |
| **ZM** | 26.1 | 32.10 | (+)18.6% | 45.910656 | (+)39.91% | 47.8236 | (+)45.37% |
| **PtLat** | 5.01 | 7.82 | (+)36.1% | 8.79449472 | (+)39.96% | 9.160932 | (+)45.42% |
| **PtMed** | 32.60 | 25.38 | (-)21.6% | 37.800576 | (+)13.75% | 39.3756 | (+)21.59% |

#### Masticatory muscles fiber length

The fiber lengths show a difference ranging from -5.23% to +8.8% (Table 5)

### Bite force data and comparisons

The comparison between the bite forces estimated in this study and those reported in the literature shows a difference ranging from -9.76% to +16.5% (Table 6). For most species, no comparison can be made due to their rarity (*Caenolestes fuliginosus*, *Dromiciops gliroides*) or the lack of study.

**Table 6 -.** Comparison table of the different methods used to estimate the fiber length (mm) of *Marmosa murina* muscles including the percentage difference between the virtual and manual methods and the different corrections.

| M1469 | Fiber length (mm) |  |  |
| --- | --- | --- | --- |
| Muscles | Length in manual dissection | Length in virtual dissection<br>VPDE method | % Difference |
| Masseter | 4.606 | 4.621 | (+)0.3% |
| Temporal | 5.472 | 5.625 | (+)2.8% |
| ZM | 5.18 | 5.564 | (+)7.4% |
| PtLat | 2.043 | 2.024 | (-)0.9% |
| PtMed | 2.109 | 2.028 | (-)3.8% |

## Discussion

### Comparison of virtual and manual methods for calculating PCSA

The use of virtual methods as a replacement for manual dissections is a growing topic of interest (Kaktze et al., 2022; Ginot et Blanke, 2024). Replacing manual dissection (a destructive method) with virtual dissection (a non-destructive method for the studied muscles) would help preserve rare or historical specimens. During this study, virtual methods were essential for estimating the PCSA of rare species such as *Caenolestes fuliginosus*.

Regarding the PCSA calculation (Table 2 and 3), virtual methods yielded varying results. The “virtual physiological data estimating method” or VPDE method, based on virtual dissection data (fiber lengths and volumes), showed a margin of error that stayed under 15,5% even for the bigger muscles. This method appears to be quite reliable and seems to give similar results as the ones obtained with other virtual methods for PCSA estimations (Santana, 2018; Katzke et al., 2022). The “slicing muscle method” or SM method produced consistent results for three of the five muscles (*M. masseter*, *M. temporalis*) but overestimated or underestimate the other (*M. zygomaticomandibularis* and *M. pterygoideus lateralis, M. pterygoideus medialis*) between 30% and 45%. The observed difference for both *M. pterygoideus lateralis* and *M. pterygoideus medialis* could be attributed to their small size, making virtual section visualization challenging due to insufficient resolution. The discrepancy for *M. zygomaticomandibularis* could be explained by the exclusion of the pennation. Regarding the muscle volume (ethanol and PMA corrected, without void, VPDE method; Figure 4 and 5), the margin of error ranges from -33.01 to +53.97%. The greatest differences are found in the *M. pterygoideus lateralis* and *M. pterygoideus medialis*, which are the smallest muscles. Consequently, the margin of error is likely to be size-dependent. When only looking at the difference in terms of relative area, the variation remains under 7.22 µm², which must have least influence as the *M. pterygoideus lateralis* and *M. pterygoideus medialis* are less implicated in the bite force generation.

Nevertheless, muscle volume remains a topic of debate since some studies report similar volumes between manual and virtual dissections (Katzke et al., 2022), while others find significant differences depending on the method that was used (Ginot and Blanke, 2024). In our study, the variation remained under 20% between the manually and virtually obtained volumes since the muscles were not treated as cylinders, and gaps between fibers were excluded. However, eliminating gaps is challenging due to the texture variation in the masticatory muscles. Indeed, within one same muscle, such as the *M. masseter*, the texture may vary significantly, with less compact areas showing visible gaps between fibers and compact areas where gaps were harder to identify. Between distinct muscles, the structure difference (more or less void between the fibers) was even more pronounced, such as between the compact *M. temporalis* and the less compact *M. zygomaticomandibularis*. This texture variation likely affected the calculated volumes. Additionally, the structure difference between distinct muscles might indicate differences in muscle density. If density differences are confirmed, PCSA calculations based on manual dissection data should also account for these differences by applying different muscle density.

Comparing results from different methods for obtaining the components of PCSA (muscle volume and fiber length) revealed different margins of error. For the fiber length the error was acceptable and ranged between -5.23% and +8.8% (VPDE method; Table 6 and 7). In this study, fiber length was simply estimated from direct measurements on CT scan images (conventional or synchrotron). However, virtual fiber lengths can be more precisely estimated, with minimal error (Katzke et al., 2022; Ginot and Blanke, 2024) using various techniques, such as importing and processing landmarks in R, using software like Amira (ThermoFisher Scientific) or ImageXd, the technique proposed by Püffel et al. (2021), or the R package *GoodFibes* (Arbour, 2023). In the future, it may be worthwhile to use one of these methods to validate the data obtained in this study. Finally, all error margins were calculated for only two species and a single individual per species, so they do not reflect intraspecific variation. These are preliminary results that require more specimens in order to obtain more accurate estimations.

**Table 7-.** Comparison table of the different methods used to estimate the fiber length (mm) of *Monodelphis touan* muscles including the percentage difference between the virtual and manual methods and the different corrections.

| M2838 | Fiber length (mm) |  |  |
| --- | --- | --- | --- |
| Muscles | Length in manual dissection | Length in virtual dissection<br>VPDE method | % Difference |
| Masseter | 6.08 | 5.85 | (-)3.99% |
| Temporal | 5.67 | 5.39 | (-)5.23% |
| ZM | 5.28 | 5.62 | (+)6.01% |
| PtLat | 2.69 | 2.81 | (+)4.32% |
| PtMed | 3.023 | 3.31 | (+)8.8% |

### Variability in modern marsupials

This study highlights significant variability in the morphology of masticatory muscles among modern marsupials, particularly concerning attachment areas, fiber lengths, masses, and contributions to total muscle mass. These findings, though based on a limited number of specimens, support the variability observed in Abreu and Astùa (2025). Furthermore, here we calculate theoretical bite force calculation on species other than the Virginia opossum, *Didelphis virginiana*, which is frequently used as a reference model for marsupials in the literature (Turnbull, 1970; Diogo et al., 2016). We describe the morphology of the masticatory muscles in some species that were not previously documented, such as *Monodelphis touan* (based on manual dissection data and virtual dissection data), *Dromiciops gliroides* and *Caenolestes fuliginosus* (using only virtual dissection data). However, these new descriptions must be interpreted cautiously. Virtual dissections are less precise than manual ones, as they were performed on entire muscles without separating their individual fascicles. This assessment followed the virtual dissection of *Marmosa murina*, where the fascicles of the *M. temporalis* could not be separated (Decuypere et al. in press). Virtual dissection, therefore, requires prior knowledge of muscle and skeletal anatomy for greater accuracy. Additionally, tendons may not always be accounted for during virtual dissection. This was the case for the tendon connecting the *M. masseter superficialis* to its origin in *Marmosa murina*, which was not visible during virtual dissection, reducing the attachment area of the muscle’s origin and impacting the centroid position of its origin area.

The preliminary data that we here provide already show variation in the position of the muscular attachments of the masticatory muscles in marsupials. This is particularly obvious when we look at the position of the insertion of the *M. zygomaticomandibularis*, whose insertion area is the largest in *Monodelphis touan* and the smallest in *Caenolestes fuliginosus*. These differences in the positioning of the attachment areas have an impact on the position of the centroid, on the resulting lever arm and therefore on the final bite force. In addition to the position of the attachment areas, muscles differ in their individual percentage of the total weight of the masticatory apparatus (see Appendix 2). These differences could be related to different muscle involvement in different species due to the constraints of diets, as has already been suggested on a larger scale (Ercoli et al., 2023). It is important to point out that these descriptions followed a nomenclature that does not match that of other authors such as Astùa and Abreu (2025), explaining at least in part the differences found between these descriptions. In fact, no specific nomenclature has been clearly defined for marsupials, although an attempt of homogenization has been made by Druzinsky et al. (2011). This represents a major problem that complicates comparisons and should be corrected in the future.

### Comparison of bite forces

The sample that we studied also showed a wide range of bite forces from the smallest specimen (belonging to the frugivore *Dromiciops gliroides)* to the largest (belonging to the carnivore *Dasyurus viverrinus)*. As previously suggested, marsupial bite force could be related to diet, although this has been refuted at the scale of Didelphidae (Abreu and Astùa, 2025). So far, our study cannot answer this debate, as our sample is too small and the effect of evolutive allometry could not be tested. This effect remains to be studied at a larger scale. The bite forces we have calculated (Table 8) are consistent with bite force estimates based on biomechanical models in the literature (Wroe et al., 2005; Abreu and Astùa, 2025), suggesting that the biomechanical model we have used can be considered as reliable. Nonetheless, it is important to note that our theoretical force calculations are based on two constants that have so far only been calculated on a small sample of domesticated placentals (Mendez and Keys, 1960; Nigg and Herzog, 1994; Leonard et al., 2020). However, jaw opening is quite different between placentals and marsupials, with a greater jaw opening potential for the latter (Attard et al., 2011; Wroe et al., 2013), suggesting a different muscular extension ability. This could be related to the constants used in the theoretical force calculation, indicating that one (or both) of them may be wrong for marsupials, as mentioned in Decuypere et al (in press). These constants should be recalculated in the future to ensure the accuracy of our theoretical model.

**Table 8:**
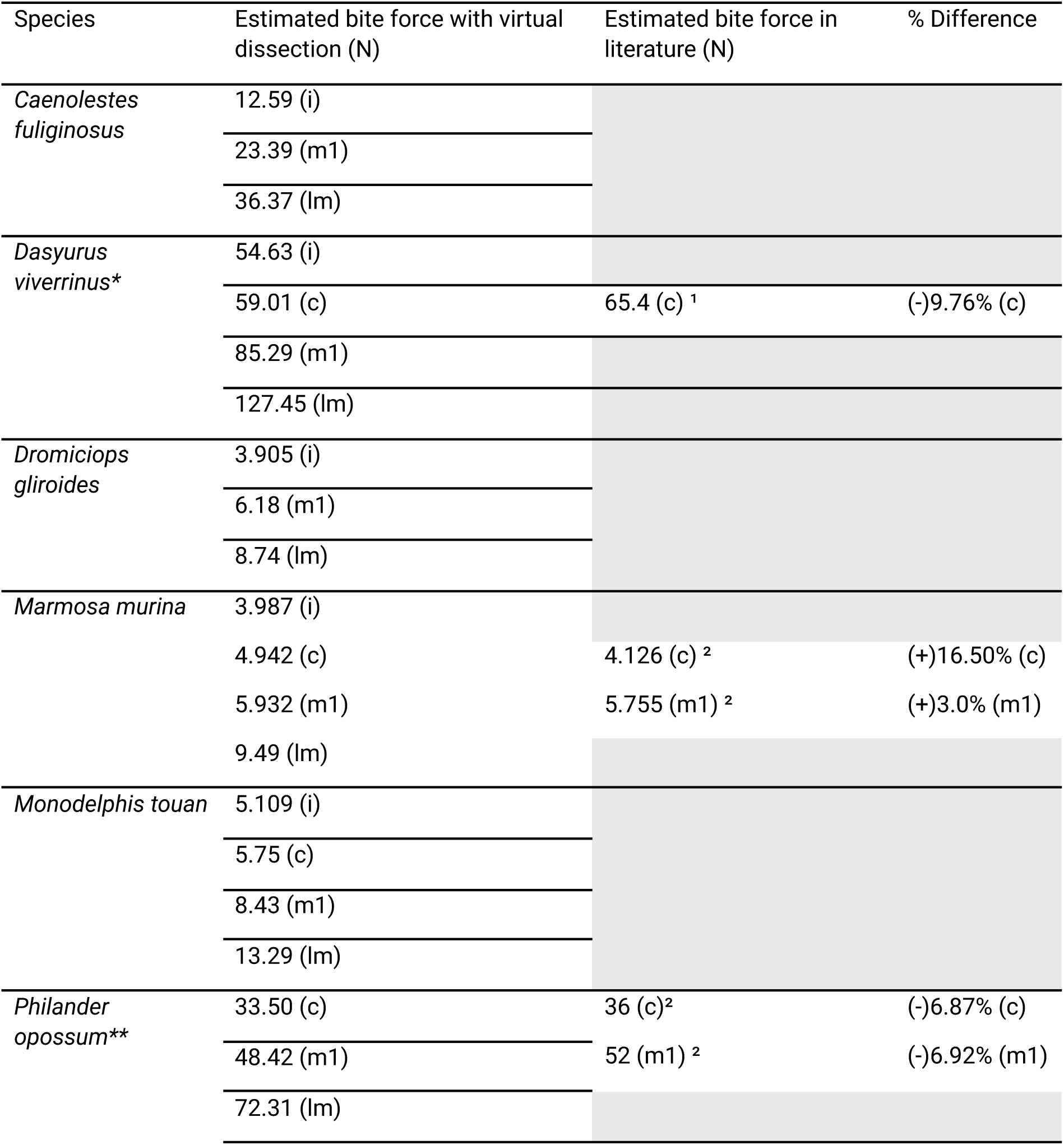
Comparison of the bite forces estimated in this study and the forces estimated in the literature. Percentage differences were calculated to compare the variation between the different results. **Abbreviations:** (c): canines, (i): incisors, (lm): last lower molar, (m1): first lower molar. Muscular informations are based from *Thomas et al. (2024) and **Abreu and Astùa (2025) Estimate bite force in literature are ¹: Based on data from Wroe et al. 2005. ²: Based on data from Abreu et Astùa (2025).

## Appendices

**Appendix 1:**
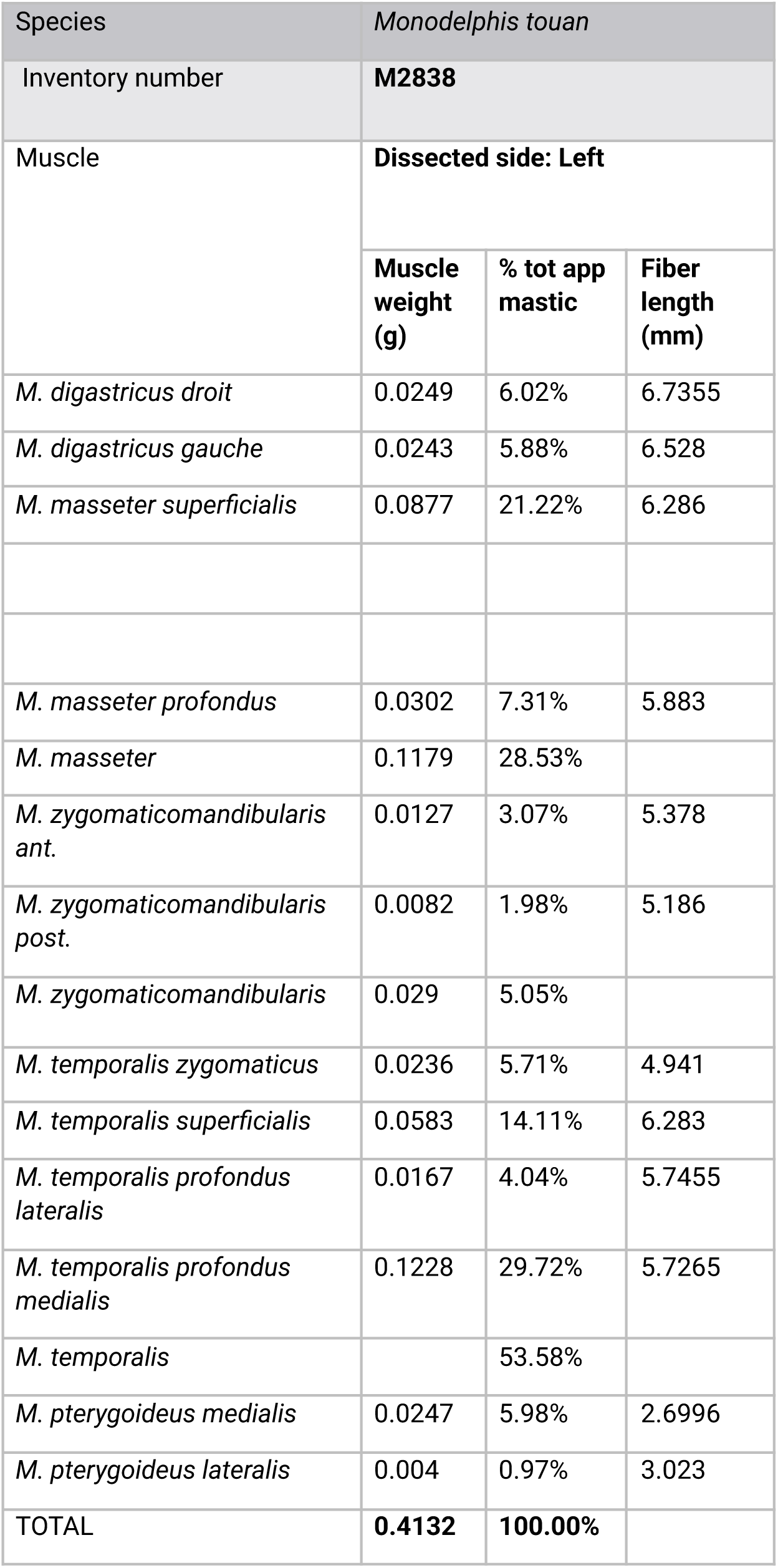
Data extracted from manual dissections of Monodelphis touan M2838.

**Appendix 2:**
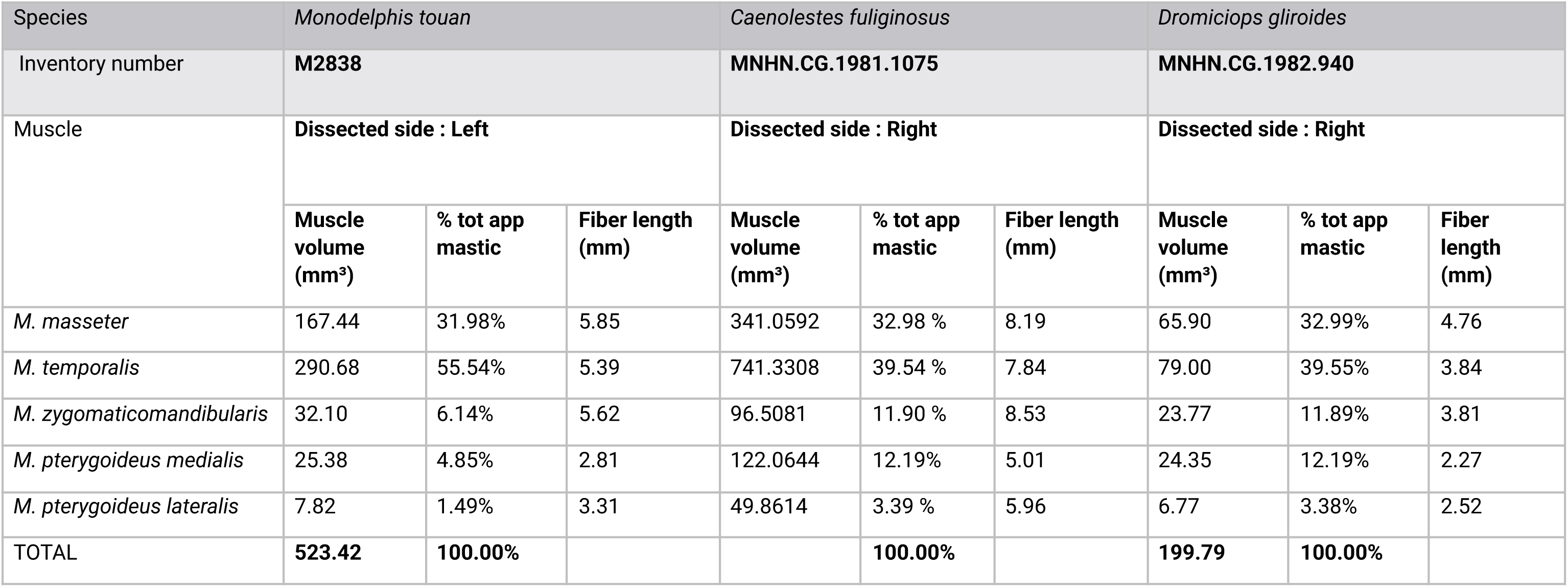
Data extracted from virtual dissections of Caenolestes fuliginosus, Dromiciops gliroides and Monodelphis touan.

## Acknowledgements

We are thankful to Benoît de Thoisy (Institut Pasteur de Guyane and KWATA association) and Anne-Claire Fabre (Institute of Ecology & Evolution, Universität Bern, Switzerland) for the collect and the use of *Marmosa murina* (M1496) and *Monodelphis touan* (M2838). We also wish to thank M. Bellato, the operator of AST-RX (MNHN), for the CT acquisitions as well as N. Poulet and F. Goussard from the 3D platforms of the CR2P for their valuable insights on segmentation and all their helpful suggestions. Our gratitude is extended to Diego Astùa and Juann A. F. H. Abreu (Laboratorio de Mastozoologia, Universidade Federal de Pernambuco, Brazil) for sharing their data and contributing to productive discussions.

## Data, scripts, code, and supplementary information availability

Raw data (CT image) will soon be available online: https://www.indores.fr/.

