## Supplemental Table 1 & 2 for "Anatomical description of the jaw muscles and theoretical bite force assessment in South-American opossums using manual and virtual dissection methods"

### Appendices

**Appendix 1:** Data extracted from manual dissections of *Monodelphis touan* M2838.

| Species | *Monodelphis touan* | | |
|---|---|---|---|
| Inventory number | **M2838** | | |
| Muscle | **Dissected side: Left** | | |
| | **Muscle weight (g)** | **% tot app mastic** | **Fiber length (mm)** |
| *M. digastricus droit* | 0.0249 | 6.02% | 6.7355 |
| *M. digastricus gauche* | 0.0243 | 5.88% | 6.528 |
| *M. masseter superficialis* | 0.0877 | 21.22% | 6.286 |
| *M. masseter profondus* | 0.0302 | 7.31% | 5.883 |
| *M. masseter* | 0.1179 | 28.53% | |
| *M. zygomaticomandibularis ant.* | 0.0127 | 3.07% | 5.378 |
| *M. zygomaticomandibularis post.* | 0.0082 | 1.98% | 5.186 |
| *M. zygomaticomandibularis* | 0.029 | 5.05% | |
| *M. temporalis zygomaticus* | 0.0236 | 5.71% | 4.941 |
| *M. temporalis superficialis* | 0.0583 | 14.11% | 6.283 |
| *M. temporalis profondus lateralis* | 0.0167 | 4.04% | 5.7455 |
| *M. temporalis profondus medialis* | 0.1228 | 29.72% | 5.7265 |
| *M. temporalis* | | 53.58% | |
| *M. pterygoideus medialis* | 0.0247 | 5.98% | 2.6996 |
| *M. pterygoideus lateralis* | 0.004 | 0.97% | 3.023 |
| TOTAL | **0.4132** | **100.00%** | |

**Appendix 2:** Data extracted from virtual dissections of *Caenolestes fuliginosus*, *Dromiciops gliroides* and *Monodelphis touan*.

| Species | *Monodelphis touan* | | | *Caenolestes fuliginosus* | | | *Dromiciops gliroides* | | |
|---|---|---|---|---|---|---|---|---|---|
| Inventory number | **M2838** | | | **MNHN.CG.1981.1075** | | | **MNHN.CG.1982.940** | | |
| Muscle | **Dissected side : Left** | | | **Dissected side : Right** | | | **Dissected side : Right** | | |
| | **Muscle volume (mm³)** | **% tot app mastic** | **Fiber length (mm)** | **Muscle volume (mm³)** | **% tot app mastic** | **Fiber length (mm)** | **Muscle volume (mm³)** | **% tot app mastic** | **Fiber length (mm)** |
| *M. masseter* | 167.44 | 31.98% | 5.85 | 341.0592 | 32.98 % | 8.19 | 65.90 | 32.99% | 4.76 |
| *M. temporalis* | 290.68 | 55.54% | 5.39 | 741.3308 | 39.54 % | 7.84 | 79.00 | 39.55% | 3.84 |
| *M. zygomaticomandibularis* | 32.10 | 6.14% | 5.62 | 96.5081 | 11.90 % | 8.53 | 23.77 | 11.89% | 3.81 |
| *M. pterygoideus medialis* | 25.38 | 4.85% | 2.81 | 122.0644 | 12.19% | 5.01 | 24.35 | 12.19% | 2.27 |
| *M. pterygoideus lateralis* | 7.82 | 1.49% | 3.31 | 49.8614 | 3.39 % | 5.96 | 6.77 | 3.38% | 2.52 |
| TOTAL | **523.42** | **100.00%** | | | **100.00 %** | | **199.79** | **100.00 %** | |
